# Accurate detection of m6A RNA modifications in native RNA sequences

**DOI:** 10.1101/525741

**Authors:** Huanle Liu, Oguzhan Begik, Morghan C Lucas, Christopher E. Mason, Schraga Schwartz, John S. Mattick, Martin A. Smith, Eva Maria Novoa

**Author notes:** Present address: Green templeton College, Oxford OX2 6HG, United Kingdom (John S. Mattick). Correspondence to: Eva Maria Novoa.

## Abstract

The field of epitranscriptomics has undergone an enormous expansion in the last few years; however, a major limitation is the lack of generic methods to map RNA modifications transcriptome-wide. Here we show that using Oxford Nanopore Technologies, N6-methyladenosine (m6A) RNA modifications can be detected with high accuracy, in the form of systematic errors and decreased base-calling qualities. Our results open new avenues to investigate the universe of RNA modifications with single nucleotide resolution, in individual RNA molecules.

## BACKGROUND

In the last few years, our ability to map RNA modifications transcriptome-wide has revolutionized our understanding of how these chemical entities shape cellular processes, modulate cancer risk, and govern cellular fate [1–4]. Systematic efforts to characterize this regulatory layer have revealed that RNA modifications are far more widespread than previously thought, can be subjected to dynamic regulation, and can profoundly impact RNA processing stability and translation [5–10]. A fundamental challenge in the field, however, is the lack of a generic approach for mapping and quantifying RNA modifications, as well as the lack of single molecule resolution [11].

Current technologies to map the epitranscriptome rely on next-generation sequencing and, as such, they are typically blind to nucleotide modifications. Consequently, indirect methods are required to identify RNA modifications transcriptome-wide, which has been mainly approached using two different strategies: (i) *antibody immunoprecipitation*, which specifically recognizes the modified ribonucleotide [5,6,12–14]; and (ii) *chemical-based detection*, using chemical compounds that selectively react with the modified ribonucleotide of interest, followed by reverse-transcription of the RNA fragment, which leads to accumulation of reads that have the same identical ends [8,9,15]. Although these methods have provided highly valuable information, they are limited by the available repertoire of commercial antibodies and the lack of selective chemical reactivities towards a particular RNA modification [16], often lack single nucleotide resolution [5–7] or require complex protocols to achieve it [17], cannot provide quantitative estimates of the stoichiometry of the modification at a given site, and are often unable to identify the underlying RNA molecule that is modified.

To overcome these limitations, third-generation sequencing technologies, such as the platforms provided by Oxford Nanopore Technologies (ONT) [18] and Pacific Biosciences (PacBio) ([3], have been proposed as a new means to detect RNA modifications in native RNA sequences. These technologies can detect RNA modifications by measuring the kinetics of the reverse transcriptase as it encounters a modified RNA -in the case of PacBio-, or by directly sequencing the RNA in its native form by pulling a native RNA through the nanopore -in the case of ONT-. Although ONT direct RNA sequencing is already a reality [19,20], extracting RNA modification information from ONT reads is an unsolved challenge. RNA modifications are known to cause disruptions in the pore current that can be detected upon comparison of raw current intensities — also known as ‘squiggles’— [18,19]. However, current efforts have not yet yielded an efficient and accurate RNA modification detection algorithm, largely due to the challenges in the alignment and re-squiggling of RNA current intensities.

As an alternative strategy, we hypothesized that the current intensity changes caused by the presence of RNA modifications may lead to increased ‘errors’ and decreased qualities from the output of base-calling algorithms that do not model base modifications (**Figure 1A**). Indeed, here we find that base-calling ‘errors’ can accurately identify N6-methyladenosine (m6A) RNA modifications in native RNA sequences, and propose a novel algorithm, *EpiNano* (github.com/enovoa/EpiNano), which can be used to identify m6A RNA modifications from native RNA reads with an overall accuracy of ~90%. Our results provide a proof of concept for the use of base-called features to identify RNA modifications using direct RNA sequencing, and open new avenues to explore additional RNA modifications in the future.

**Figure 1.**
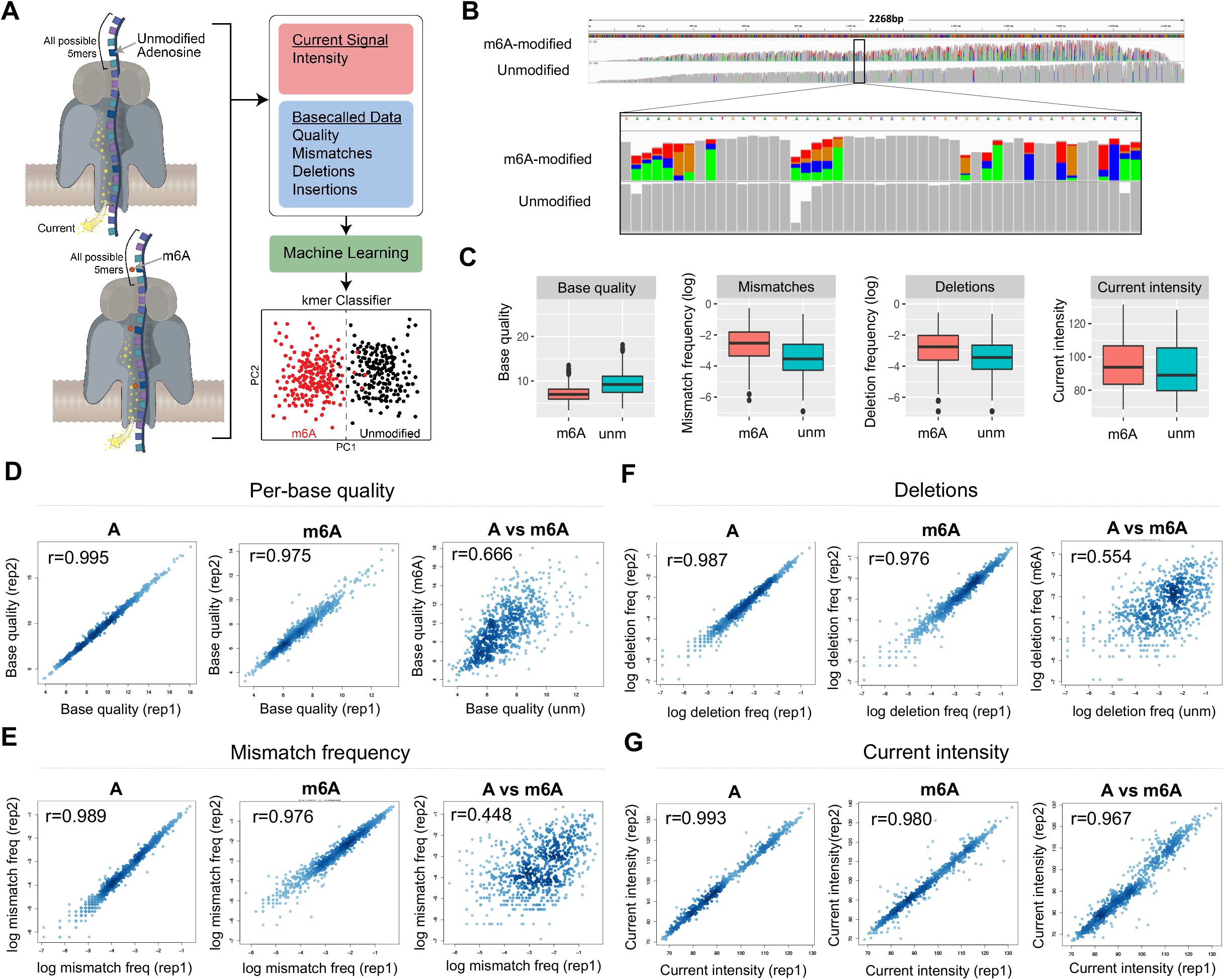
Base-calling ‘errors’ can be used as a proxy to identify RNA modifications in direct RNA sequencing reads. (**A**) Schematic overview of the strategy used in this work to train and test an m6A RNA base-calling algorithm (**B**) IGV snapshot of one of the four transcripts used in this work. In the upper panel, *in-vitro* transcribed products containing m6A have been mapped, whereas in the lower panel the unmodified counterpart is shown. Nucleotides with mismatch frequencies greater than 0.05 have been coloured. **(C)** Comparison of m6A and A positions, at the level of per-base quality scores (left panel), mismatch frequencies (middle left panel), deletion frequency (middle right panel) and mean current intensity (right panel). All possible k-mers (computed as a sliding window along the transcripts) have been included for these comparisons (n=9974) (**D, E, F, G**) Replicability of each individual feature - base quality (D), deletion frequency (E), mismatch frequency (F) and current intensity (G)-across biological replicates, for both unmodified (‘A’) and m6A-modified (‘m6A’) datasets. Comparison of unmodified and m6A-modified (‘A vs m6A’) is also shown for each feature. Correlation values shown correspond to Spearman’s rho.

## RESULTS AND DISCUSSION

Previous work has shown that ONT raw current intensity signals, known as ‘squiggles’, can be subdivided into ‘events’, which correspond to consecutive 5-mer sequences shifted one nucleotide at a time (e.g., in the sequence AGACAAU, the corresponding 5-mer ‘events’ would be AGACA, GACAA, and ACAAU) [21–24]. Therefore, to systematically identify the current intensity changes caused by the presence of a given RNA modification, perturbations of the current intensity signals should be measured and analyzed for each possible 5-mer (n=1024). To this end, we designed a set of synthetic sequences that comprised all possible 5-mers (median occurrence of each 5-mer=10), while minimizing the RNA secondary structure (see Methods and **File S1**). We then compared the direct RNA sequencing reads of *in* vitro-transcribed constructs that incorporated N6-methyladenosine (‘m6A’) to those with unmodified ribonucleotides (‘unm’) (**Figure 1A**). Comparison of the two datasets revealed that base-called m6A-modified reads are significantly enriched in mismatches compared to their unmodified counterparts (**Figure 1B and 1C**), and that these ‘errors’ are mainly, but not exclusively, located in adenine positions. We observed that, in addition to mismatch frequency, other metrics including per-base quality, insertion frequency, deletion frequency and current intensity, were significantly altered (**Figure 1C** and **Figure S1**). Moreover, these ‘errors’ were reproducible in independent biological replicates with respect to mismatch frequency, deletion frequency, per-base quality and current intensity (**Figures 1D-G**). By contrast, insertion frequencies were not reproducible across biological replicates (**Figure S1**), suggesting that this feature is likely unrelated to the presence of RNA modifications, and thus was not further considered in downstream analyses.

We then examined whether these observed differences would be sufficient to accurately classify a given site into “modified” or “unmodified”. For this aim, we first focused our analysis on 5-mers that matched the known m6A motif RRACH, as these would be the most relevant in which to detect m6A modifications. To reveal whether the features from m6A-modified RRACH k-mers were distinct from unmodified RRACH k-mers, we compiled the base-called features (base quality, mismatch frequency and deletion frequency) for each position of the k-mer (−2, −1, 0, +1, +2) (**Figure 2A**, see also **Figure S2**), and performed Principal Component Analysis (PCA) of the features, finding that the two populations (m6A-modified and unmodified RRACH k-mers) were largely non-overlapping (**Figure 2B**). As a control, we performed the same analysis in k-mers with identical sequence context, but centered in C, G, or U (instead of A), finding that no differences could be observed between these populations (**Figure 2C**), suggesting that the observed differences are m6A-specific and not dataset-specific.

**Figure 2.**
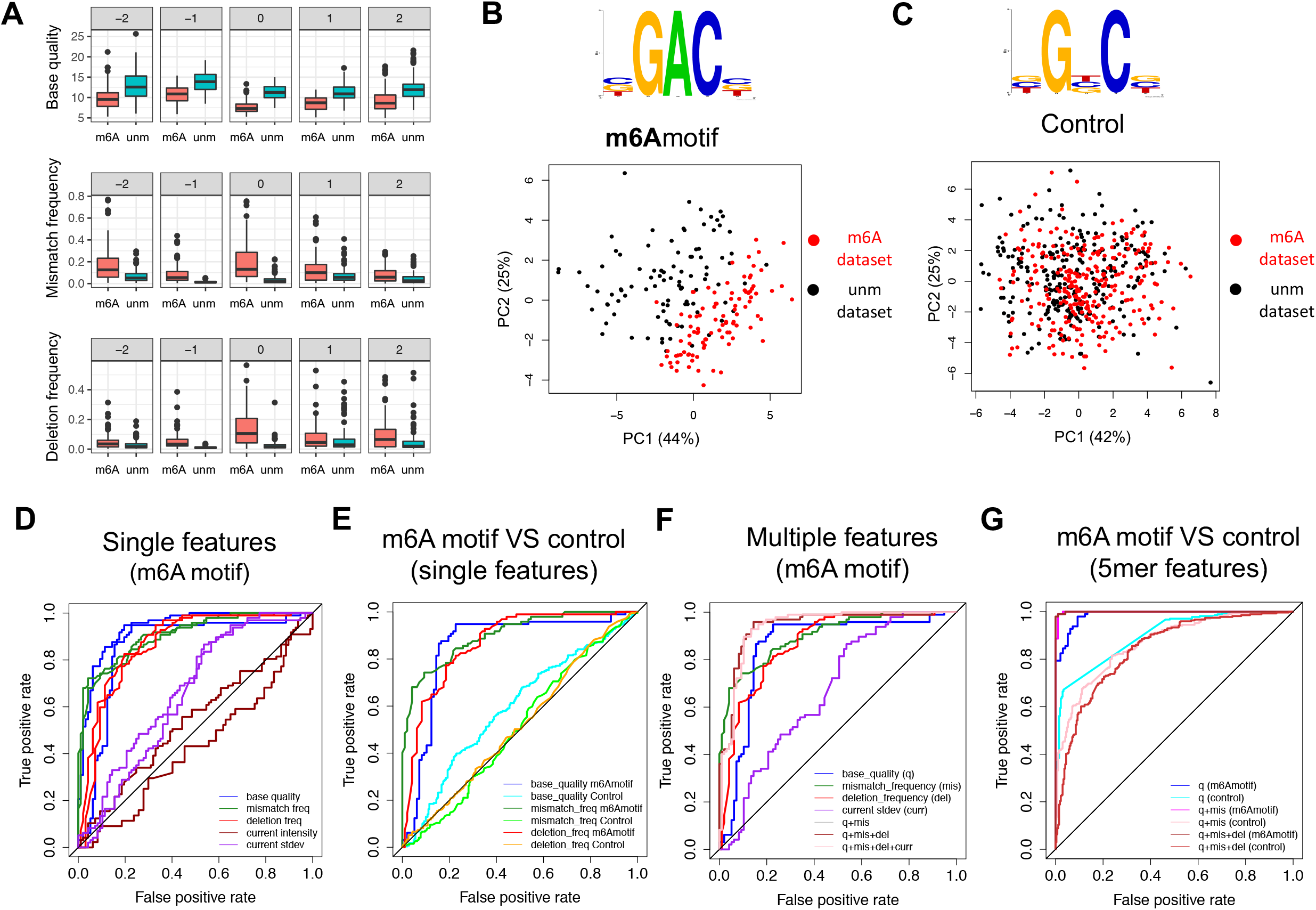
Base-calling ‘errors’ alone can accurately identify m6A RNA modifications. (**A**) Base-called features (base quality, insertion frequency and deletion frequency) of m6A motif 5-mers, and for each position of the 5-mer, are shown. The features of the m6A-modified transcripts (‘m6A’) are shown in red, whereas the features of the unmodified transcripts (‘unm’) are shown in blue. (**B, C**) Principal component analysis (PCA) scores plot of the two first principal components, using 15 features (base quality, mismatch frequency, deletion frequency, for each of the 5 positions of the k-mer) as input. The logos of the k-mers used in the m6A-motif RRACH set (left) and control set (right) are also shown. Each dot represents a specific k-mer in the synthetic sequence, and has been coloured depending on whether the k-mer belongs to the m6A-modified transcripts (red) or the unmodified transcripts (black). The contribution of each principal component is shown in each axis. (**D, E, F, G**) ROC curves of the SVM predictions using: i) each individual feature separately to train and test each model, at m6A sites (D); ii) each individual feature separately, comparing m6A motif k-mers to control k-mers (E); iii) combined features at m6A sites, relative to the individual features (F); and iv) combined features at m6A sites relative to control sites, where the base-called ‘errors’ information of neighbouring nucleotides has been included in the model (G).

To statistically determine whether these features could be used to accurately classify a given site into ‘m6A-modified’ or ‘unmodified’, we trained multiple Support Vector Machines (SVM) using as input the base-called features from m6A-containing RRACH k-mers and unmodified RRACH k-mers (see Methods). We first tested whether each individual feature at position 0 (the modified site) was able to classify a given RRACH k-mer into m6A-modified or unmodified. Our results show that base quality, deletion frequency and mismatch frequency alone were able to accurately predict the modification status with reasonable accuracy (70-86% accuracy, depending on the feature used) (**Figure 2D**, see also **Table S1** and Methods). By contrast, we find that the current mean intensity values and current intensity standard deviation were poor predictors of the modification status of the k-mer (43-65% accuracy). As a control, we used the same set of features in control k-mers (i.e., those with the same sequence context, but centered in C, G or U), finding that the features did not distinguish between m6A-modified and m6A-unmodified datasets (**Figure 2E**, see also **Figure S3**). To improve the performance of the algorithm, we then examined whether a combination of the features might improve the prediction accuracy, finding that the combination of the 3 features (base quality, mismatch and deletion frequency) increased the accuracy of the model (88-91%) (**Figure 2F,** see also **Table S1**). Finally, we tested whether the inclusion of all the features from the neighbouring positions (−2, −1, +1, +2) would improve the performance of the algorithm. We find that the inclusion of neighbouring features slightly improves the performance of the algorithm (accuracy = 97-99%), however, this was at the expense of increasing the detection of false positives that in the control k-mer set -which do not contain the modification- (**Figure 2G,** see also **Figure S3**), suggesting that features from neighbouring positions should not be employed with this model.

Overall, our results provide a proof of principle for the use of base-calling ‘errors’ as an accurate and computationally simple solution to identify m6A modifications with an overall accuracy of ~90%. Moreover, our method does not require the manipulation of raw current intensities or squiggle alignments. We envision that similar approaches can be developed to detect other RNA modifications in the future, opening new avenues to explore the role of RNA modifications for which we currently lack transcriptome-wide maps. In addition, our m6A-modified and unmodified datasets covering all possible 5-mers, which are publicly available, can be employed to train different machine learning algorithms than the ones tested here (e.g., signal-based machine learning, base-caller training, etc.), and thus obtain improved methods to detect RNA modifications in the future.

## CONCLUSIONS

The human epitranscriptome is still largely uncharted. Only a handful of the 170 different RNA modifications that are known to exist have been mapped. Importantly, several of these modifications have been found to be involved in central biological processes, such as sex determination or cell fate transition, and their dysregulation has been linked to multiple human diseases, including neurological disorders and cancers. Yet, our understanding of this regulatory layer is restricted to a few RNA modifications, largely due to the lack of a generic methodology to map them in a transcriptome-wide fashion. This work provides a novel strategy to identify RNA modifications from base-called features, without the need of squiggle realignments, opening new avenues to study the epitranscriptome with unprecedented resolution. The establishment of the ONT platform as a tool to map virtually any given modification will allow us to query the epitranscriptome in ways that, until now, had not been possible. Future work can expand to other modifications like 5-methylcytosine (m5C), as well as provide additional thresholds for controlling specificity and sensitivity.

## ADDITIONAL FILES

### Supplemental materials and Methods

**File S1.** Fasta sequences of the in vitro constructs used in this work to train and test the RNA modification algorithm.

**File S2.** Step-by-step protocol to produce m6A-modified and unmodified RNA sequences, to be sequenced using direct RNA sequencing, and used to train/test the RNA modification detection algorithm.

### Supplementary figures

**Figure S1.**
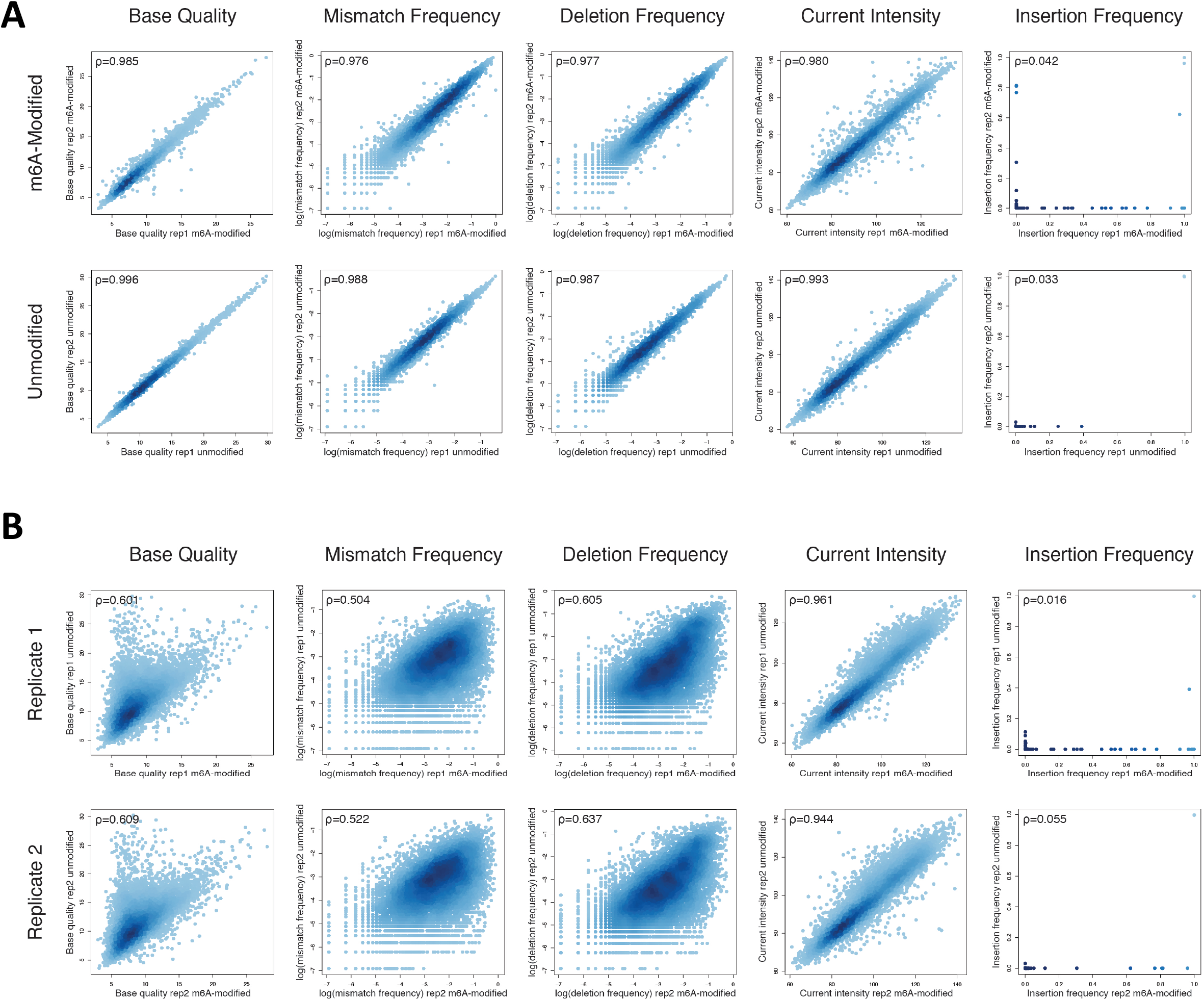
Replicability of the features extracted. Each dot corresponds to a different nucleotide of the synthetic constructs (n=9978)

**Figure S2.**
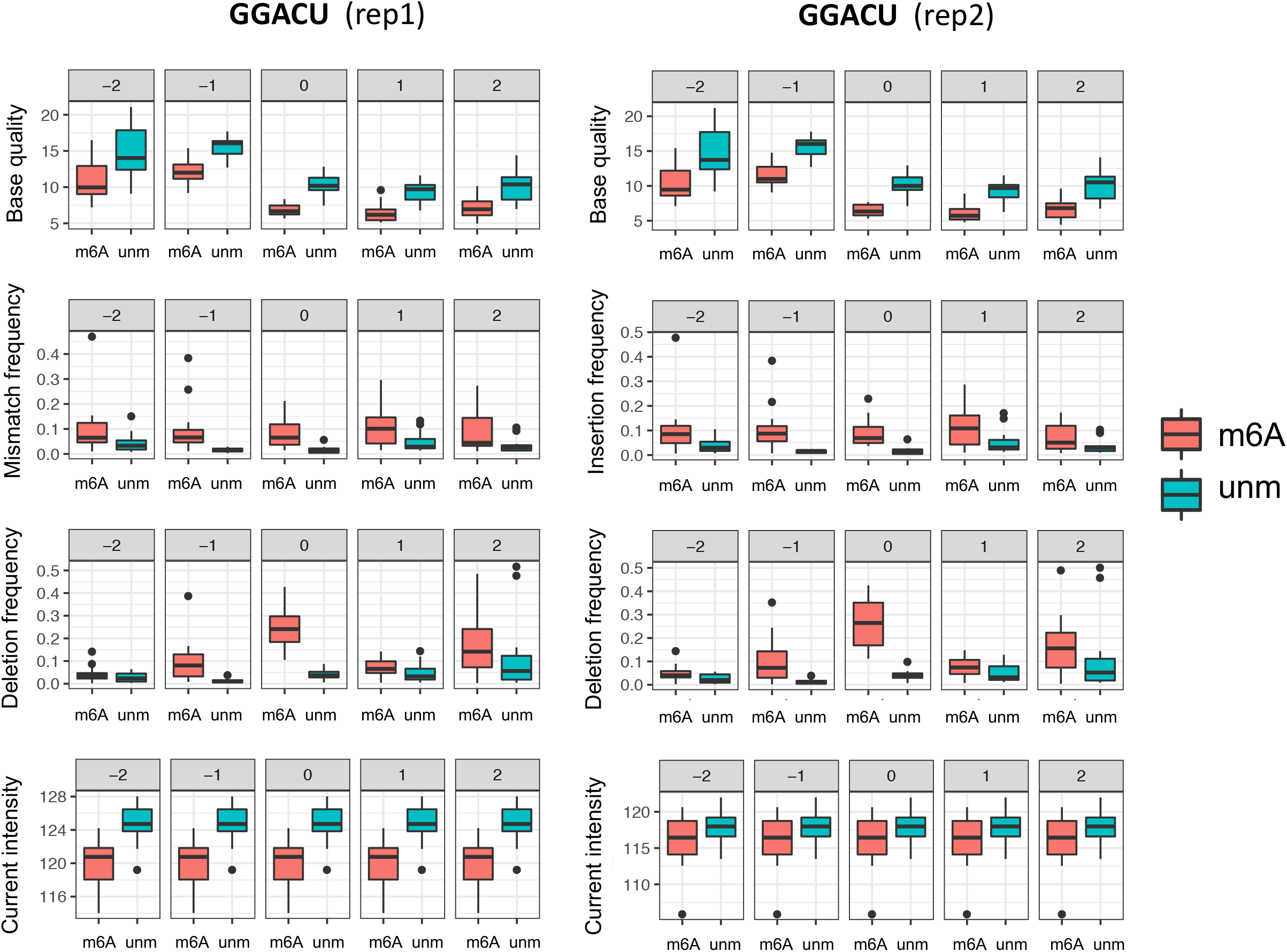
Replicability of the base-called features of GGACU k-mers, for each position of the k-mers. Base-called features of m6A-modified datasets are depicted in red, whereas those from unmodified datasets are depicted in blue.

**Figure S3.**
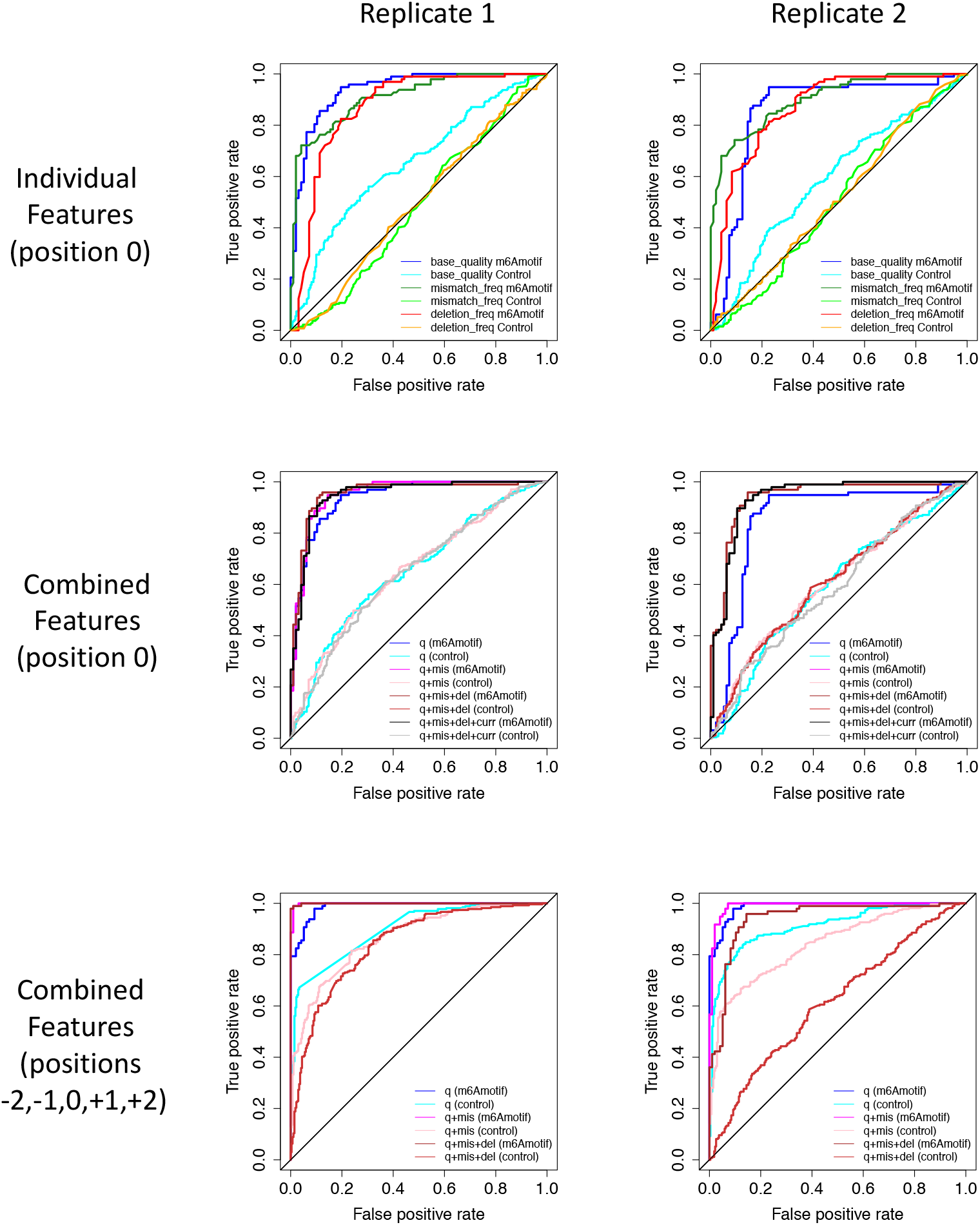
ROC curves of SVM trained with single features compared to combined features. Performance of each replicate is shown separately in each plot.

**Figure S4.**
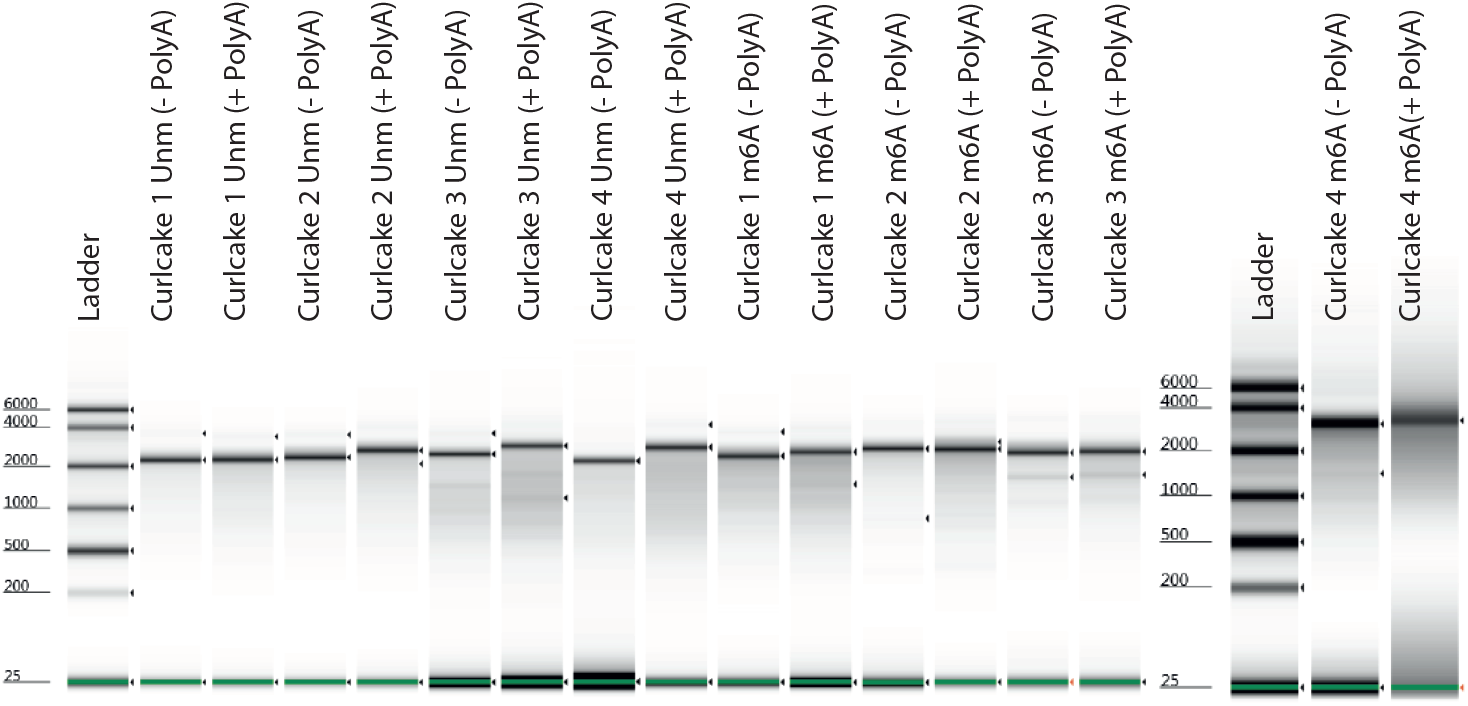
TapeStation output of the quality and quantity of the m6A-modified and unmodified *in vitro* transcribed products

### Supplementary tables

**Table S1.** Accuracy of prediction of m6A-modified sites in m6A motifs, relative to control k-mers, using different feature combinations, and for both replicates.

**Table S2.** Metrics of quality and yield of *in vitro* transcription of both modified and unmodified RNAs, which were used as input for the direct RNA sequencing runs.

**Table S3.** Metrics of the number of sequenced reads, base-called reads and mapped reads using direct RNA sequencing, for the 4 runs included in this study

## METHODS

### Synthetic sequence design

Sequences were designed such that they would include all possible 5-mers, while minimizing the secondary RNA structure. For this aim, we employed the software *curlcake* (http://cb.csail.mit.edu/cb/curlcake/), which internally uses RNAshapes version 2.1.6 [25] to predict RNA secondary structure. The final output sequence given by the software was ~10kb long. For synthesis purposes, a total of 4 sequences were designed by splitting the 10kb sequence into smaller sequences of slightly different size (2329bp, 2543bp, 2678bp and 2795bp, which we named ‘Curlcake 1’, ‘Curlcake 2’, ‘Curlcake 3’ and ‘Curlcake 4’, respectively). Each sequence was designed with an internal strong T7 polymerase promoter, an additional BamHI site at the end of the sequence, and with all EcoRV and BamHI sites removed from the sequence (**File S1**). All 4 sequences were synthesized and cloned in pUC57 vector using blunt EcoRV by General Biosystems. Plasmids were double digested O/N with EcoRV-BamHI-HF, and DNA was extracted with Phenol-Chloroform followed by EtOH precipitation. Plasmid digestion was confirmed by agarose gel. Digestion product quality was assessed with Nanodrop before proceeding to *in vitro* transcription (IVT).

### In vitro transcription, capping and polyadenylation

*In vitro* transcribed (IVT) sequences were produced using the Ampliscribe™ T7-Flash™ Transcription Kit (Lucigen-ASF3507), using 1 ug of purified digestion product as starting material, following manufacturer’s recommendations. ATP was replaced by N^6^-Methyladenosine-5’-Triphosphate(m^6^ATP) (Trilink-N-1013;) for the IVT reaction of m6A-modified RNA. IVT reaction was incubated for 4 hours at 42°C. *In vitro* transcribed RNA was then incubated with DNAse I (Lucigen), followed by purification using RNeasy Mini Kit (Qiagen-74104). Integrity and quality of the RNA was determined using Agilent 4200 Tapestation, to ensure that a single product band of the correct size had been produced for each IVT product (**Figure S4**). Each IVT product was 5’ capped using Vaccinia Capping Enzyme (NEB-M2080S) following manufacturer’s recommendations. The capping reaction was incubated for 30 minutes at 37 °C. Capped IVT products were purified using RNA Clean XP Beads (Beckman Coulter-A66514). Poly(A)-tailing was performed using *E. coli* Poly(A) Polymerase kit (NEB-M0276S), following manufacturer’s recommendations. Poly(A)-tailed RNAs were purified using RNA Clean XP beads, and the addition of poly(A)-tail was confirmed using Agilent 4200 Tapestation (**Figure S4**). Concentration was determined using Qubit Fluorometric Quantitation. Purity of the IVT product was measured with NanoDrop 2000 Spectrophotometer (**Table S2**)

### Direct RNA library preparation

RNA library for direct RNA Sequencing (SQK-RNA001) was prepared following the ONT Direct RNA Sequencing protocol version DRS_9026_v1_revP_15Dec2016. Briefly, 800 nanograms of Poly(A)-tailed and capped RNA — 200 nanograms per construct-was ligated to ONT RT Adaptor (RTA) using concentrated T4 DNA Ligase (NEB-M0202T), and was reverse transcribed using SuperScript III Reverse Transcriptase (Thermo Fisher Scientific-18080044). The products were purified using 1.8X Agencourt RNAClean XP beads (Fisher Scientific-NC0068576), washing with 70% freshly prepared ethanol. RNA Adapter (RMX) was ligated onto the RNA:DNA hybrid, and the mix was purified using 1X Agencourt RNAClean XP beads, washing with Wash buffer (WSB) twice. The sample was then eluted in Elution Buffer (ELB) and mixed with RNA running buffer (RRB) prior to loading onto a primed R9.4.1 flow cell, and ran on a GridION (MinION for the second replicate) sequencer with MinKNOW acquisition software version v1.14.1 (v. 1.15.1 for the second replicate). The sequencing was performed in independent days and machines, with two biological replicates for each condition (non-modified and m6A-modified RNA).

### Base-calling, filtering and mapping

Reads were locally base-called using Albacore 2.1.7 (Oxford Nanopore Technologies). Base-called reads were filtered using NanoFilt, a component from Nanopack [26], with settings ‘-q 0 --headcrop 5 --tailcrop 3’, and mapped to the 4 synthetic sequences using minimap2 [27] with the settings -ax map-ont. Mapped reads were then converted into mpileup format using Samtools version 1.4. [28]. Statistics on read basecalling and mapping can be found in **Table S3**.

### Feature extraction

To extract per-site features (mean per-base quality, mismatch frequency, insertion frequency and deletion frequency), BAM alignment files were converted to tab delimited format using sam2tsv from javakit [29]. For each individual reference site, the mean quality of the aligned bases, the mismatch, insertion and deletion frequency was computed using in-house scripts (available on github). To extract current intensity information from individual reads, the h5py (version 2.7.0) module in python was used to parse each individual fast5 file. Reference sequences were slided with a window size of 5bp, and mean and standard deviation of current intensities was computed for each sliding window. All in-house python scripts used to extract the features described above are publicly available as part of *EpiNano* (github.com/enovoa/EpiNano).

### Machine learning

The set of extracted features was used as input to train a Support Vector Machine (SVM), where 75% of the sites were used for training, whereas 25% of the sites were used for testing, using five-fold cross validation. Multiple kernels (‘linear’, ‘poly’ and ‘rbf’) were compared. Each trained SVM was validated on a new run of freshly *in vitro* transcribed m6A-modified and unmodified sequences, which had not been used for initial training or testing of the SVM. The code to generate the set of features for machine learning, the code for building the SVM models, as well as the trained SVM models are publicly available in github (github.com/enovoa/EpiNano). We should note that a limitation in utilizing *in vitro* transcription to generate all possible 5-mers is that 5-mers that contain more than one “A” will contain more than one modification in the kmer, e.g. AGACC will in fact be m6AGm6ACC, which are unlikely to occur in a biological context. Therefore, 5-mers that contained more than one A have been excluded from the analyses. Accuracy of the model has been computed as the sum of correct m6A modification predictions - correctly predicted m6A-modified k-mers (true positives, TP) and correctly predicted unmodified k-mers (true negatives, TN)- divided by the total number of k-mers tested. Five-fold cross-validation was used to define the

## LIST OF ABBREVIATIONS

(m6A): N6-methlyadenosine
(m1A): N1-methyladenosine
(5hmc): 5-hydroxymethylcytosine
(ONT): Oxford Nanopore Technologies
(SVM): Support Vector Machine
(IVT): *In vitro* transcription

## DECLARATIONS

### Availability of data and material

All code used in this work is publicly available at github.com/enovoa/EpiNano. All FASTQ files data generated in this work have been made publicly available at the GEO database under the accession code GSE124309.

### Funding and acknowledgements

O.B. is supported by an international PhD fellowship (UIPA) from the University of New South Wales. MCL is supported by CRG International PhD Fellowships Programme. E.M.N was supported by a DECRA fellowship from the Australian Research Council (DE170100506) and is currently supported by CRG Severo Ochoa Funding. This work was funded by the Australian Research Council (DP180103571). We acknowledge support of the Spanish Ministry of Economy, Industry and Competitiveness (MEIC) to the EMBL partnership, Centro de Excelencia Severo Ochoa and CERCA Programme / Generalitat de Catalunya. C.E.M thanks funding from the Bert L and N Kuggie Vallee Foundation, the WorldQuant Foundation, The Pershing Square Sohn Cancer Research Alliance, NASA (NNX14AH50G, NNX17AB26G), the National Institutes of Health (R01ES021006, 1R21AI129851, 1R01MH117406), the Bill and Melinda Gates Foundation (OPP1151054), the Leukemia and Lymphoma Society grants (LLS 9238-16, LLS-MCL-982). We would like to thank James Ferguson for all his helpful comments, as well as for giving us early access to his fast5 processing toolkit (https://github.com/Psy-Fer/fast5_fetcher).

### Author’s contributions

HL performed the bioinformatics analysis, together with EMN. OB and MCL performed the experimental work including the preparation and running of the direct RNA sequencing libraries. HL, OB, MCL and EMN prepared the figures. EMN and MAS conceived the project. EMN supervised the project, with contribution of SS, CEM, JM and MAS. HL, OB, MCL and EMN wrote the manuscript, with the contribution of all authors.

